# Forest condition in the Congo Basin for the assessment of ecosystem conservation status

**DOI:** 10.1101/2020.03.25.008110

**Authors:** Aurélie Shapiro, Hedley S. Grantham, Naikoa Aguilar-Amuchastegui, Nicholas J. Murray, Valery Gond, Djoan Bonfils, Olivia Rickenbach

## Abstract

Measuring forest degradation is important for understanding and designing measures to protect biodiversity and the capacity of forests to deliver ecosystem services. Conservation planning, particularly the prioritization of management interventions for forests, is often lacking spatial data on ecological condition, and it is often overlooked within decision-making processes. Existing methods for assessing forest degradation via proxies or binary measures (i.e. intact or not) cannot adequately consider the important variations of ecological condition. Direct methods to measure degradation (e.g. through remote sensing) require extensive training data, calibration and validation, and may be too sensitive to small-scale or short-term changes which may not be related to degradation. We developed a metric termed Forest Condition (FC) which aims to measure the degree of forest degradation, incorporating temporal history of forest change over a large spatial extent. We parameterized this metric based on estimated changes in above ground biomass in the context of forest fragmentation over time to estimate a continuous measure of forest degradation for Congo Basin countries. We estimate that just less than 70% of Congo Basin forests remain fully intact. FC was validated by direct remote sensing measurements from Landsat imagery for DRC. Results showed that FC was significantly positively correlated with forest canopy cover, gap area per hectare, and magnitude of temporal change in Normalized Burn Ratio. We tested the ability of FC to distinguish primary and secondary degradation and deforestation and found significant differences in gap area and spectral anomalies to validate our theoretical model. We used the IUCN Red List of Ecosystems criteria to demonstrate the value of applying forest degradation to assess the risk of ecosystem collapse. Based on this assessment, we found that without including FC in the assessment of biotic disruption, 12 ecosystems could not have a threat status assigned, and a further 9 ecosystems would have a lower threat status. Our overall assessment of ecosystems found approximately half of forest of Congo Basin ecosystem types which cover over 20% of all forest area are threatened including 4 ecosystems (<1% of total area) which are critically engendered. FC is a transferrable and scalable assessment to support forest monitoring, planning, and management.

## 1. Introduction

Forests provide valuable ecosystem services to people, such as provision of food and materials, hydrological functions for clean supply of water, and home to numerous indigenous peoples (Díaz et al., 2019). Forests are also at the forefront of global initiatives for the mitigation of greenhouse gas emissions, as conserving remaining intact forests is important for carbon sequestration and avoidance of future potential emissions (Jantz et al., 2014; Mitchell et al., 2017). Forests harbour unique and important biodiversity which underpins many of these ecosystem functions, aligning with conservation efforts (Feeley and Terborgh, 2005; Stokstad, 2014). Intact forest ecosystems are shown to have greater conservation benefits than degraded or fragmented ones of similar ecological type (Betts and et al., 2019; Haddad et al., 2015), making strong arguments for prioritizing them for conservation management (Watson et al., 2018).

Forests are increasingly threatened by expanding human activities (Thompson et al., 2011; Venter et al., 2016). This results in the degradation of forests through a process of fragmentation, which in turn impacts biodiversity, biomass, and therefore the ability of forest to provide many ecosystem services (Betts and et al., 2019; Chaplin-Kramer et al., 2015; Haddad et al., 2015; Potapov et al., 2012). Although there is no standard definition of forest degradation (Ghazoul et al., 2015; Potapov et al., 2009), it has been acknowledged that declines in forest intactness result in environmental and social problems which impact forest health, affecting human livelihoods and economic development (Foley et al., 2005; Pereira et al., 2010). Understanding and quantifying changes in forest fragmentation related to ecological condition is therefore crucial to monitor, manage and protect more intact forests over time to prevent such problems (Brooks et al., 2006; Mittermeier et al., 2003). We define degradation via the term forest “condition” using a combination of spatial patterns of fragmentation and ecosystem services, notably above ground biomass (AGB) as described in Shapiro et al., 2016.

Remote sensing can provide affordable, efficient, consistent multi-temporal measurements for forest monitoring, and determination of forest condition when appropriately defined (Mitchell et al., 2017). The recent increases in the reliable use of satellite constellations, as well as improved access to data and enhanced processing capabilities, are promoting analyses of higher temporal resolution which enable improved assessments of forest degradation over time. Remote sensing approaches for forest degradation are generally grouped into direct and indirect approaches (Herold et al., 2011). There are advantages and disadvantages to each approach which will vary by geography, resources available, and specific needs. Direct remote sensing methods estimate parameters or spectral indices related to canopy gaps and structure, changes in forest canopies, or productivity in time series (DeVries et al., 2015; Mitchell et al., 2017; Souza et al., 2005; Spruce et al., 2011; Verbesselt et al., 2012, 2010), although the implementation over a large area is limited by image resolution or availability of time series or consistency between sensor types or climate effects (Cohen et al., 2010; Kennedy et al., 2010; Zhu, 2017) which hinders the ability to compare variables in different geographies or climate regimes. Direct satellite measurements are also affected by the complexity of defining degradation according to specific remote sensing indicators, and are more sensitive to forest dynamics, changes in vegetation, climate or even extreme events such as droughts, which may represent shorter term events which may be confused with degradation. In contrast, indirect methods employ the mapping of proxies, for example presence of roads, fires, forest edges or pattern (Broadbent et al., 2008; Chaplin-Kramer et al., 2015; Haddad et al., 2015; Potapov et al., 2008; Riitters et al., 2015; Shapiro et al., 2016; Tyukavina et al., 2016). These methods are particularly suitable for planning and monitoring, reporting and verification in developing countries with low field monitoring resources (Bucki et al., 2012). Fragmentation and spatial pattern approaches are conceptually simpler, and being used in the development of reference levels and targets for emissions reduction programs, for example in Nepal (Forest Carbon Partnership Facility (FCPF), 2018).

A simple approach for assessing and monitoring forest condition can provide a repeatable, transferrable and understandable indicator for effective regional conservation planning and prioritization for conservation. An indicator that provides a continuous estimation forest condition can provide richer information than binary variables (for example intact forest landscapes) which need to be defined by thresholds specific to certain geographies.

In this study, we build on previous research (Shapiro et al., 2016), to assess forest condition (FC) by developing analyses of key forest fragmentation and structure indicators over time. We first assess changes in forest spatial pattern, and then use available estimates of above-ground biomass (AGB) in strata defined by these spatial patterns to assign a continuous estimation of FC. FC is calculated by effects of fragmentation and degradation of forest edges over time using relative changes in AGB. We apply a theoretical model to discern primary and secondary degradation from deforestation, and demonstrate how the results -- a new forest condition metric – enable evaluations of the extent and severity of ecosystem degradation to assess ecosystem collapse under the IUCN Red List of Ecosystems categories and criteria (Bland et al., 2015; IUCN, 2016a; Rodríguez et al., 2015)

## 2. Materials and Methods

### 2.1 Study Area

The Congo Basin forest ecoregion is comprised of tropical forests in the Democratic Republic of Congo (DRC), Republic of Congo (ROC), Equatorial Guinea, Gabon, Cameroon, Central Africa Republic and a small portion of Angola (Figure 1). This represents the largest tract of forest in Africa, and the single largest peatland complex in the world, storing a significant amount of forest carbon (Dargie et al., 2017). The basin is highly biodiverse and is a focus of recent species discovery (Dargie et al., 2019; Hart et al., 2012), while more than 30 million people inhabit the basin, including indigenous communities with a long and intricate relationship to natural ecosystems (Riddell, 2013). Together these characteristics represent a unique ecological opportunity to mitigate climate, while supporting the livelihoods of the many communities who depend on essential natural resources. The relative lack of information, compared to the Amazon basin or Asian forests currently hinders successful management and conservation efforts in the context of needed sustainable development (Malhi et al., 2013).

**Figure 1:**
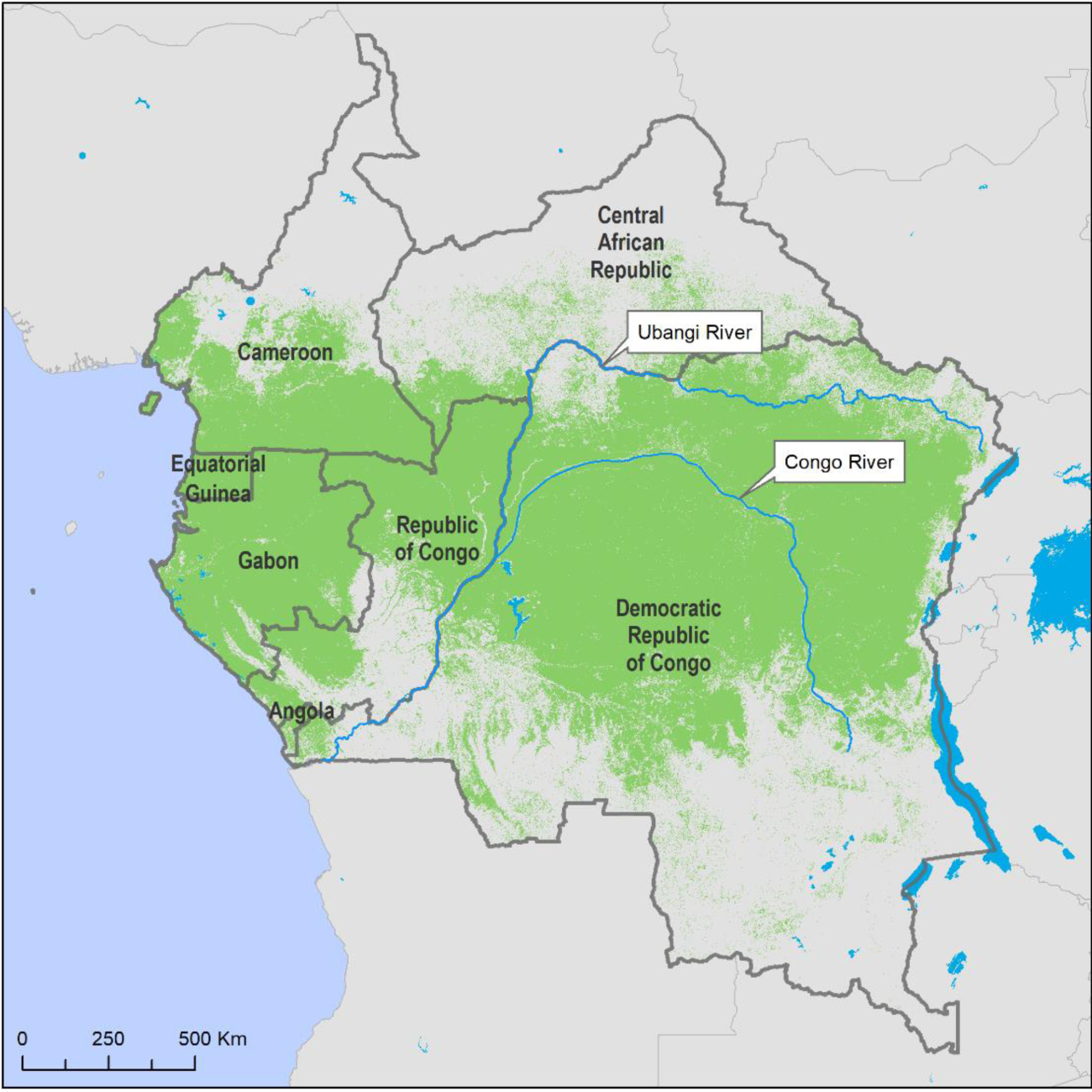
The regional study area encompasses 6 countries in the Congo Basin. A national scale assessment focuses on the Democratic Republic of Congo (DRC). Major biogeographic boundaries are defined by the Ubangi and Congo Rivers. Green shows the primary tropical forest cover for 2016.

### 2.2 Data Sources

We developed a comprehensive dataset of relevant ecological, physical and forest data layers to evaluate FC for Congo Basin forests.

#### 2.2.1 Congo Basin Forest Ecosystems

We acquired forest cover data for Congo Basin countries for terra firme forests from Philippon *et al*. (2018), which use phenology patterns and light regimes derived from MODIS (Moderate Resolution Imaging Spectrometer) to identify 8 distinct forest types at 500m resolution. To complete coverage of all forest types in our study area we identified open forests using data from Hansen et al., 2013, which were identified from treecover greater than 60% (in 2000). We integrated mangroves mapped by Giri et al., 2011. Lastly, we added swamp forest types by overlaying data from two sources, Betbeder *et al*. 2014 and Dargie *et al*. 2017, which together identified 14 unique swamp forest types by flooding dynamics and dominant species (see supplemental material). We resampled our forest types data to a common pixel resolution of 1 ha (100 m x 100m).

To better represent known biogeographic patterns in forest types, we split our combined maps into bio-geographic regions known as important bio-physical barriers which have isolated distinct species of great apes for millions of years (Olson and Dinerstein, 2001; Takemoto et al., 2015). To represent these regions, we split areas east and west of the Congo River, and north and south of Ubangi river. We further distinguish sub-montane and montane vegetation according to elevations above 1100m and 1750m respectively (Verhegghen et al., 2012). Finally, we identified important Marantaceae dominated forests in the Republic of Congo (WCS, unpublished data.). The final product was a a map of 64 forest classes for the year 2000 (see supplemental material for a list of all forest ecosystem types), which was updated to 2016 by removing all areas identified as tree cover loss by Hansen et al., 2013. The forest maps for both time periods were used to create binary forest/non-forest masks.

#### 2.2.2 Above Ground Biomass (AGB)

AGB (Mg/ha) at the Congo Basin scale was sourced from the *integrated pan-tropical dataset* developed by Avitabile et al., (2016) at 1km resolution. For the local-scale assessment in DRC, we use a national dataset calibrated by airborne LiDAR (Light Detection and Ranging; Section 2.2.3) and field data, extrapolated to the national DRC scale using wall-to-wall Landsat, ALOS PALSAR active radar and topography datasets as described in Xu et al., (2018).

### 2.3 Developing a forest condition metric

We estimated FC by combining forest fragmentation change and the relative loss in AGB. First, we assigned the forest/non-forest mask from the two time periods (2000 and 2016) into fragmentation classes using Morphological Spatial Pattern Analysis (MSPA) from the GUIDOS toolbox (Soille and Vogt, 2009; Vogt and Riitters, 2017). We used an edge distance of 300m (Shapiro et al., 2016) and reclassed bridges and loops to inner and outer edges based on their location on the boundary of interior or exterior non-forest patches respectively. Thus, we classify forest in each time period into one of four fragment classes: core, inner edge, outer edge and patch forest.

We then assess transitions in fragmentation classes from 2000 to 2016. We identified areas that change from core forest to other fragmentation classes, identifying which forest pixels remain in the same class, versus transitions between different classes, which are assigned primary and secondary deforestation, primary and secondary degradation, as shown in Figure 2.

**Figure 2.**
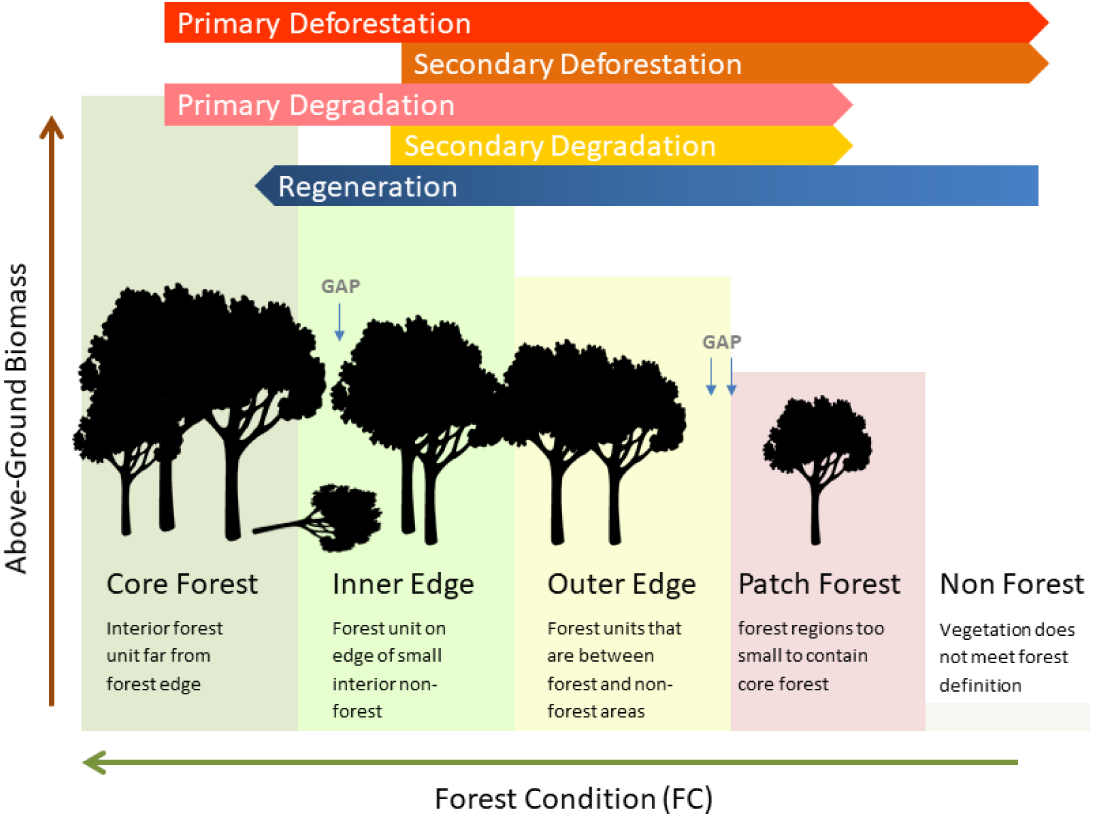
Theoretical concept of deforestation and degradation transitions via fragmentation. A forest/non-forest map is classified into 5 fragmentation types (core, inner edge, outer edge, patch forest) which have decreasing levels of above-ground biomass (AGB) respectively. The transitions between classes from one time period to the next are described in the top of the figure with an arrow that has a beginning point and an end, e.g. a change in core forest to outer edge is primary degradation. An inner edge forest that becomes non-forest is secondary deforestation. Stable forest types are primary forest (core forest with no change) and secondary forest (inner and outer edges, patch forests with no change).

The change in above-ground biomass between two time periods (2000 to 2016) was calculated from the fragmentation class transition using a process analogous to the gain-loss method for carbon stock monitoring (Murdiyarso et al., 2008). Gains and losses in AGB are calculated according to differences between fragmentation strata means (Shapiro et al., 2016).

We computed FC as a continuous metric from 0-100, based on the proportional change in biomass between classes, thereby integrating the temporal dynamics of a forest area that is an indication of not only present state (one snapshot: degraded or not) but the state in a trajectory from intact to deforested. This transition is determined according to the proportion of AGB remaining in comparison to the mean AGB of the core forest strata. Relative FC was then estimated on a continuous scale from fully intact (100) to completely lost (0), based on the proportional loss of biomass between fragmentation classes for two time periods.

FC of the second time period j for each forest ecosystem is calculated using the following Equation 1)

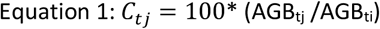

*Where C is the condition of that specific forest ecosystem fragmentation strata at any time t (denoted by tj), based on the AGB of the previous and current fragmentation category*.

To differentiate an ecosystem that has changed to a new state versus one that is stable, we assess final condition (FC) using requation (2):

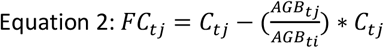

### 2.4 Validation of Forest Condition Metric in DRC

We assessed the performance of FC and the theoretical framework in several ways for the DRC, for which we have detailed validation data (Table 1Table 1. We correlated FC with fractional forest cover and canopy gap area, along with the estimate of biomass lost and the NBR cumulative anomalies. This was done using a random sample of points (50 points distributed inside the fragmentation classes inside each LiDAR plot, n=10,800, see Xu et al., 2017 for description of LiDAR data) collected for assessment with forest structure variables of fractional cover and gap area, biomass lost and anomaly using a Pearson correlation matrix executed in R software (version 3.5.1). Negative cumulative anomalies of NBR (Section 2.1.6) were evaluated within each degradation class using analysis of variance (ANOVA) of the same random sample of points as above. These were further evaluated using the Tukey honest significant difference pairwise test to determine differences between paired fragmentation transition classes.

**Table 1.** Datasets and descriptions and relevant article section

| Data | Source | Description | Section |
| --- | --- | --- | --- |
| *Data to derive FC* |  |  |  |
| Forest Ecosystems | Philippon et al., 2018<br>Hansen et al., 2013<br>Giri et al., 2011<br>Dargie et al., 2017<br>Betbeder et al., 2014 | 64 unique forest types determined by phenology, climate, flooding dynamics and bio-geographical zone | 2.2.1 |
| Above Ground Biomass | Xu et al., 2017 | National forest biomass dataset derived from LiDAR and satellite imagery for the DRC | 2.2.2 |
| *Validation Data* |  |  |  |
| Canopy height | Xu et al., 2017 | National airborne LiDAR dataset for the DRC | 2.4.1 |
| Forest gap area | Xu et al., 2017 | Derived from LiDAR canopy height following method of Betts et al., 2005 | 2.4.2 |
| Fractional cover | Xu et al., 2017 | Derived from LiDAR forest canopy height | 2.4.3 |
| Normalized Burn Ratio | Key and Benson, 2005 | Index derived from Landsat Tier 1 imagery | 2.4.4 |

#### 2.4.1 Forest canopy height

Forest canopy height was estimated using the airborne LiDAR dataset collected in 2014 and 2015 throughout the DRC following a systematic random sampling pattern, as described by the VCS VT0005 methodology (Tittmann et al., 2015; Xu et al., 2017). In total, 216 random plots of 2,000 ha each were distributed over a 1° × 1°grid laid over the national primary dense forest cover dataset for DRC (Potapov et al., 2012). LiDAR data were collected with a mean point density of 2/m^2^ from which digital surface models and mean canopy height were derived at 2m meter resolution (Xu et al., 2017). All canopy heights above 3m (national forest definition) were used to create a detailed forest cover map for these LiDAR sampling areas, and used to develop the variables described in the following two sections.

#### 2.4.2 Forest Gap Area

Forest gap area was estimated using the difference between the LIDAR canopy height and a maximum estimated within a 50-cell window (or 1 hectare, following Betts et al., 2005). Gaps were identified using a threshold of 21m less than the canopy maximum, which located all gap areas within continuous forest, verified by the very high resolution (10cm) airborne imagery collected by the same airborne data collection campaign. The gap area was then summed for each hectare in the LiDAR footprints and sampled using the random sample of 100 points per LiDAR plot.

#### 2.4.3 Fractional Cover

Forest fractional cover was estimated from the 2m forest layer by summing the total number of cells in a 50×50 window and calculating the proportion of 2500 cells covered by forest to produce % forest cover at the 1ha scale.

#### 2.4.4 Normalized Burn Ratio

We used the normalized burn ratio index (NBR; Key and Benson, 2005) as a direct remote sensing indicator of canopy disturbance associated with encroachment and illegal logging (Langner et al., 2018). We calculated NBR from Landsat surface reflectance imagery from the USGS Tier 1 collection from 1984 to 2016 processed in Google Earth Engine (Gorelick et al., 2017). All available Landsat data since 1984 were compiled, filtered by cloud cover (less than 90%), poor quality pixels were masked according to pixel quality (Foga et al., 2017), and the image collected was sorted by acquisition date. We use a cumulative anomaly analysis to assess NBR in a monitoring period (2000-2016) compared to a baseline historical period (all previously available imagery from 1984-1999), where all Landsat images are sorted in time, and the differences with the mean are sequentially summed and divided by the number of available images. From 2000 onward, coinciding with the first year of forest condition transition assessment, the difference between calculated NBR for each cloud-free pixel and the historical mean was calculated, summed, and normalized by the number of non-null observations as in Lagomasino et al., (2019). An area with a time period of high positive anomalies (higher NBR than historical mean) followed by subsequently larger negative anomalies, will have an overall high negative accumulated anomaly.

### 2.5 IUCN Redlist of Ecosystems assessment

To estimate the risk of collapse of each of the 64 Congo Basin ecosystem types, we used the IUCN Red List of Ecosystem criteria A2b, B1 and B2 and D (Bland et al., 2015). The Red List of Ecosystems is a rule-based protocol that utilises information on spatial change, range size, and biotic and abiotic variables for each ecosystem to identify ecosystems are corded according to most at risk of ecosystem collapse.

Criterion A2b was used to assess the reduction in geographic extent of each ecosystem over a 50-year period, using the adjusted proportional rate of decline based on the extent data for two time periods, 2000 and 2016 (See section 2.1.1). To assess the range size criterion B, we computed extent of occurrence as a minimum convex polygon encompassing all occurrences of each ecosystem (criterion B1) and area of occupancy using the 1% occupancy rule (criterion B2) and appropriate sub-criteria as described in Bland et al., 2015.

Criterion D focusses on the disruption of biotic processes (Bland et al 2017), for which we applied the area of primary degradation (see Figure 2) as the extent of the disruption, and the mean forest condition to indicate severity. Forest edges are known for their detrimental effects on ecosystems services and vertebrate habitats (Pfeifer et al., 2017), thus, making a fragmentation approach relevant for conservation prioritization applications. We used the year 1850 as the historical reference, as prior to then forests in the Congo Basin were considered largely free of human disturbances and industrial development (Morin-Rivat et al., 2017). Both sub-criteria D2 and D3 were evaluated to determine the validity of these assumptions.

The change in FC over the 16 year study period was used as an indicator of biotic disruption, as reduced AGB affects the delivery of ecosystem services such as climate change mitigation over time, (Heymell et al., 2011; Pettorelli et al., 2017; Shvidenko et al., 2005). The change in amount of core forest vs. edge classes determined the extent of the ecosystem affected by fragmentation, edge effects (N. M. Haddad et al., 2015) for criterion D3, while the changes in mean forest condition per ecosystem were used to assess relative severity of degradation for the severity.

FC by definition assumes that at some initial point in time, all forests were intact core ecosystems with 100% condition, thus providing the information needed to assess two of the sub-criteria D2a and D3. For D2a, we presume the rates of change of core vs. edges determine the fraction of the extent of the ecosystem affected since 2000; and these are projected to 2050 using the proportional annual rate of decline (PRD; Rodríguez *et al*. 2015). For D3, we assessed the proportional rates of decline over the actual annual rates of decline (ARD) in mean FC by ecosystem which were modelled using the changes from 1850-2016, with the assumption that in 1850, all forest ecosystems were core intact forest with maximum potential biomass (Figure 3).

**Figure 3.**
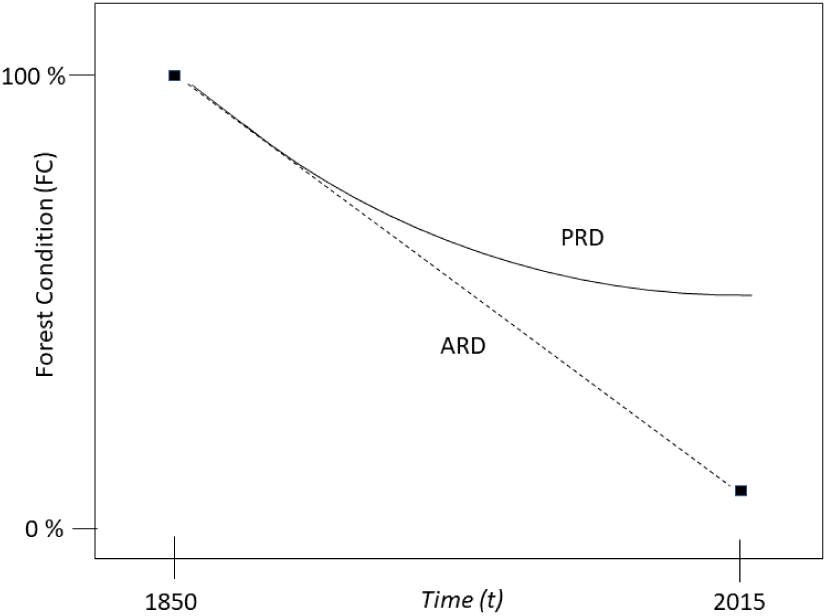
The correlation between forest condition estimated in 2015, and the assumed 100% condition in 1850, can be calculated using either ARD or PRD (adapted from Rodríguez et al., 2015).

**Figure 4.**
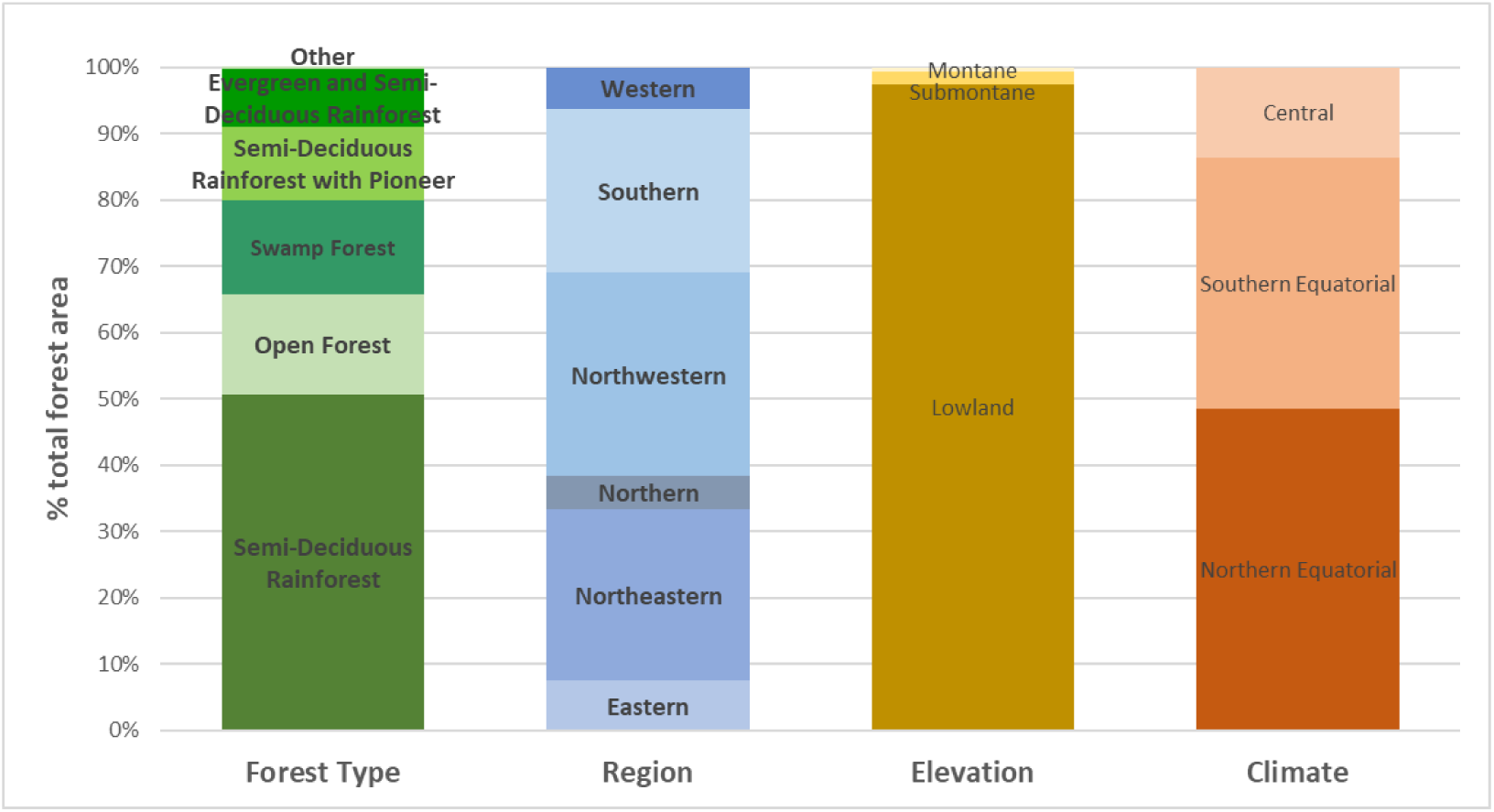
Congo Basin forest composition by region, forest type, elevation and climate. (Other forest types include mangrove and Marataceae)

The final ecosystem status was assigned as the highest assessment outcome between all three criteria evaluated, A, B and D.

## 3 Results

### 3.1 Condition of Congo Basin Ecosystems

Our forest ecosystem map shows the Congo Basin forests cover more than 210 million ha and predominantly lowland, equatorial semi-deciduous forests with a significant swamp forest ecosystem in the central region covering more than 29 million hectares.

The condition of these 64 forest ecosystems vary widely (Figure 6). For broad forest types, open forests and mangroves have the lowest mean condition, while swamp forests and the mixed evergreen and semi-deciduous rainforests have the highest mean FC (Table 2). The localized Maranthaceae forests have the highest mean condition. High condition forest (>80) is generally present in the dense forest ecosystems in Gabon, which have the highest mean forest condition, followed by Republic of Congo (Table 3). Large areas of lower condition (<50) are present in eastern DRC, along the Congo river and in the southwest corner of DRC, and south central Cameroon, while fragmented, low condition forests are predominant in the Central African Republic.

**Table 2.** Mean FC by general forest type.

| Broad Forest Type | Total Area (ha) | Mean FC | Std. dev. |
| --- | --- | --- | --- |
| Dense Evergreen Rainforest | 4,457,859 | 82.13 | 32.24 |
| Evergreen and Semi-Deciduous Rainforest | 18,177,916 | 89.92 | 25.99 |
| Semi-Deciduous Rainforest | 104,332,094 | 85.24 | 30.38 |
| Semi-Deciduous Rainforest with Pioneer | 22,453,096 | 75.14 | 38.05 |
| Maranthaceae | 267,717 | 91.71 | 20.81 |
| Swamp Forest | 28,928,944 | 85.88 | 32.02 |
| Mangrove | 402,780 | 64.71 | 40.51 |
| Open Forest | 31,239,177 | 18.93 | 15.22 |

**Table 3.** Mean FC by Congo Basin country

| Country | Total Area (ha) | Forest area 2015 (ha) | Mean FC | Std. dev. |
| --- | --- | --- | --- | --- |
| Cameroon | 47,177,546 | 21,686,790 | 75.21 | 36.41 |
| Central African Republic | 62,889,075 | 11,385,949 | 45.47 | 39.32 |
| Republic of Congo | 34,220,955 | 23,701,530 | 84.91 | 31.97 |
| Equatorial Guinea | 2,701,407 | 2,594,197 | 77.27 | 35.99 |
| Gabon | 26,489,820 | 23,939,932 | 85.94 | 29.98 |
| Democratic Republic of Congo | 234,751,788 | 126,437,088 | 73.25 | 38.58 |
| Angola | 712,269 | 514,097 | 52.46 | 43.05 |

**Figure 5.**
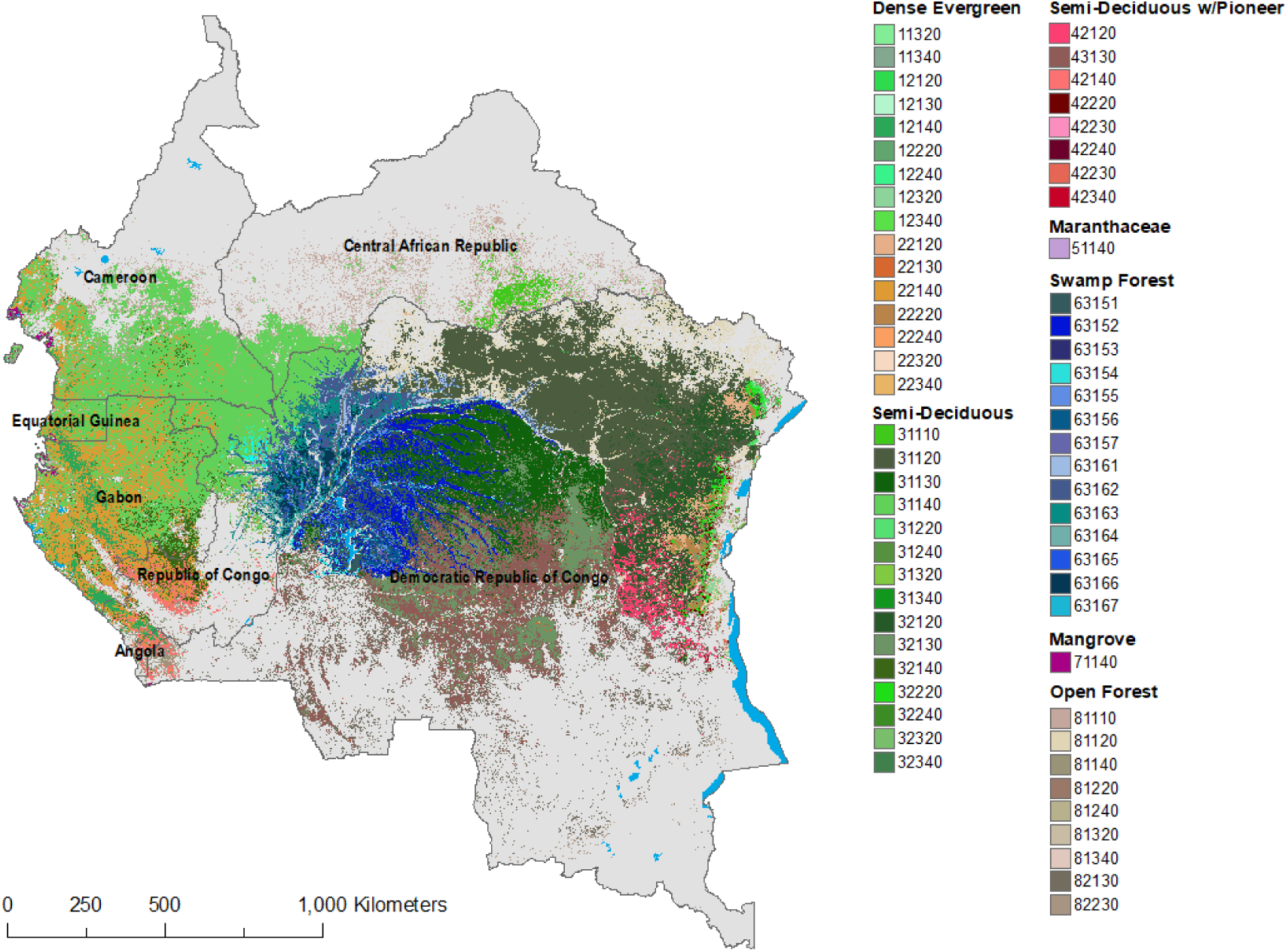
Distribution of forest ecosystem types of the Congo Basin

**Figure 6.**
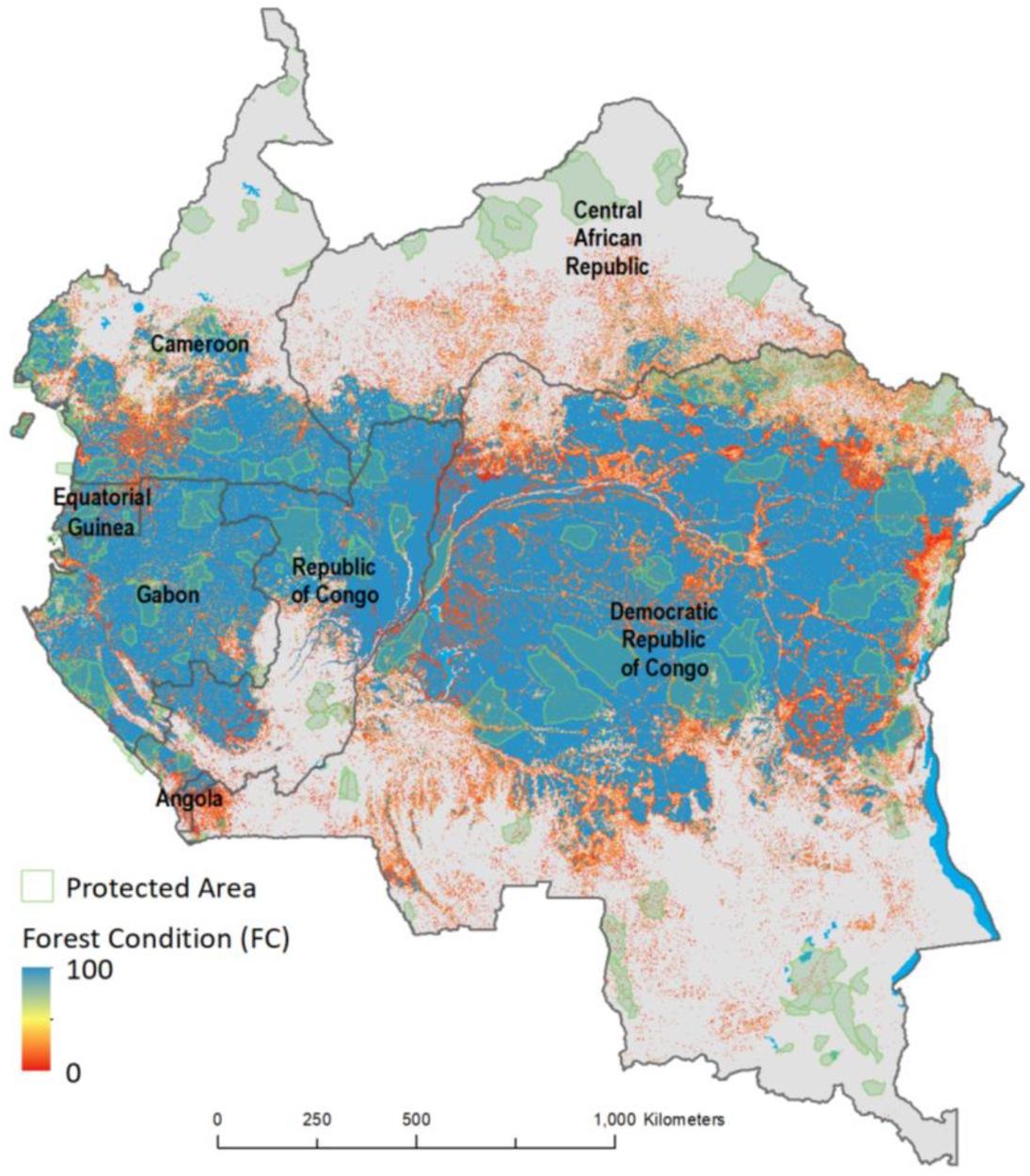
FC for Congo Basin forests (2015). Protected areas data from (Pélissier et al., 2019) and WRI Forest Atlases (World Resources Institute, 2019).

### 3.2 Validating FC in the DRC

Using the detailed LiDAR dataset for random sample plots in the DRC, (n = 21,600) FC was shown to be significantly, yet weakly correlated with fractional cover, gaps, biomass loss and NBR anomalies, with the greatest negative correlation with gaps and biomass loss (Figure 7). The NBR anomalies also show the highest positive correlation with fractional cover, where greater negative anomalies are correlated with lower fractional cover.

**Figure 7.**
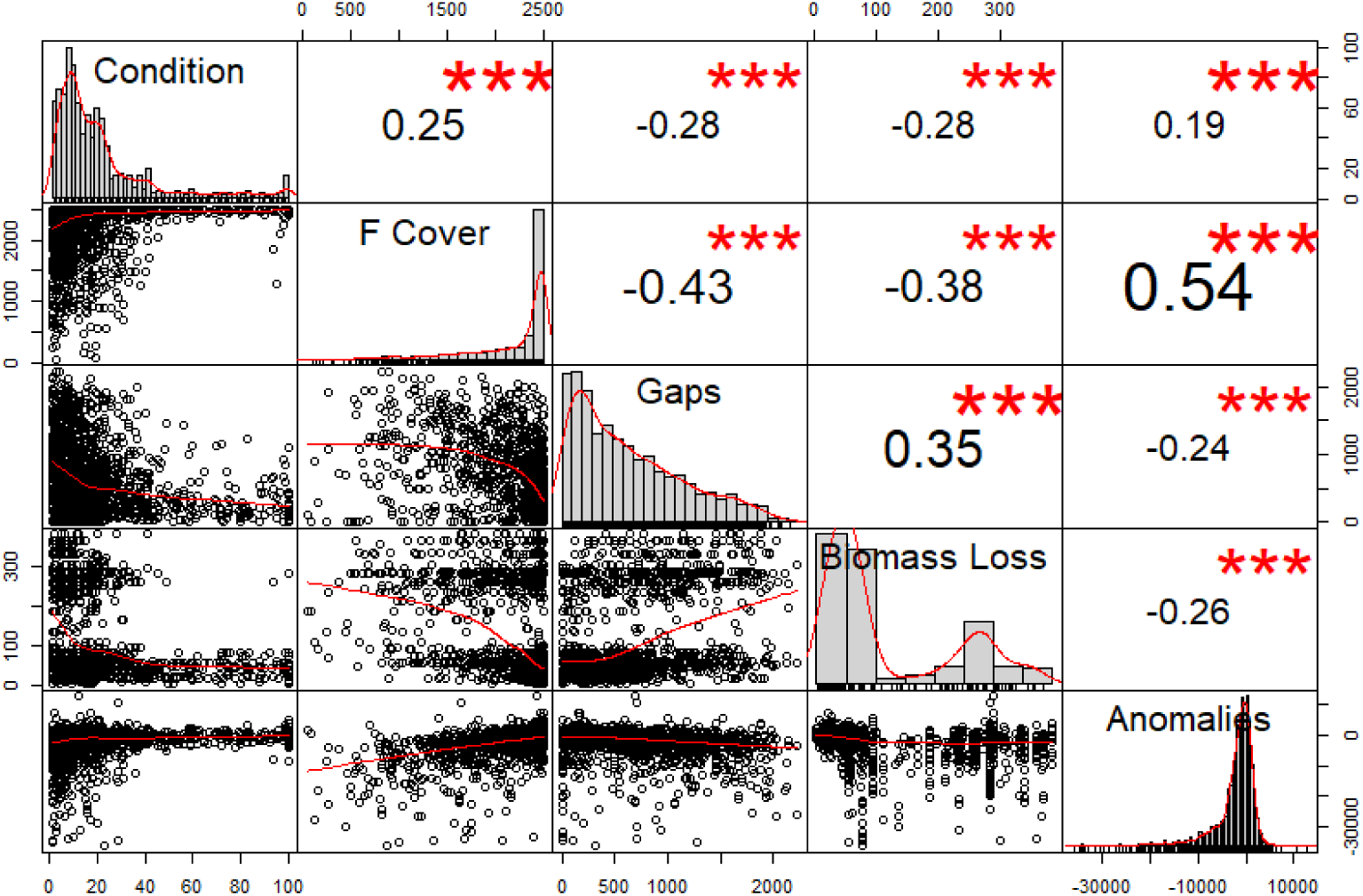
Correlation matrix of sampled variables including ecosystem condition, fractional cover at 1 ha (F Cover, Section 2.2.5), Biomass loss (Mg/ha, Section 2.2.2) and the NBR Anomalies Section (2.2.6). The distribution of each variable is shown on the diagonal, bivariate scatter plots on the lower left, and the correlation coefficient shown as a value. Significance levels are denoted by red stars (3 stars: p<0.001; 1 star: p<0.05).

When assessing gap area by transition type, gap area decreases significantly for stable classes (primary and secondary forest), with the highest gap area observed in areas which were identified as primary deforestation (Figure 8). Gap area however was not significantly different between primary and secondary deforestation and primary and secondary degradation classes.

**Figure 8.**
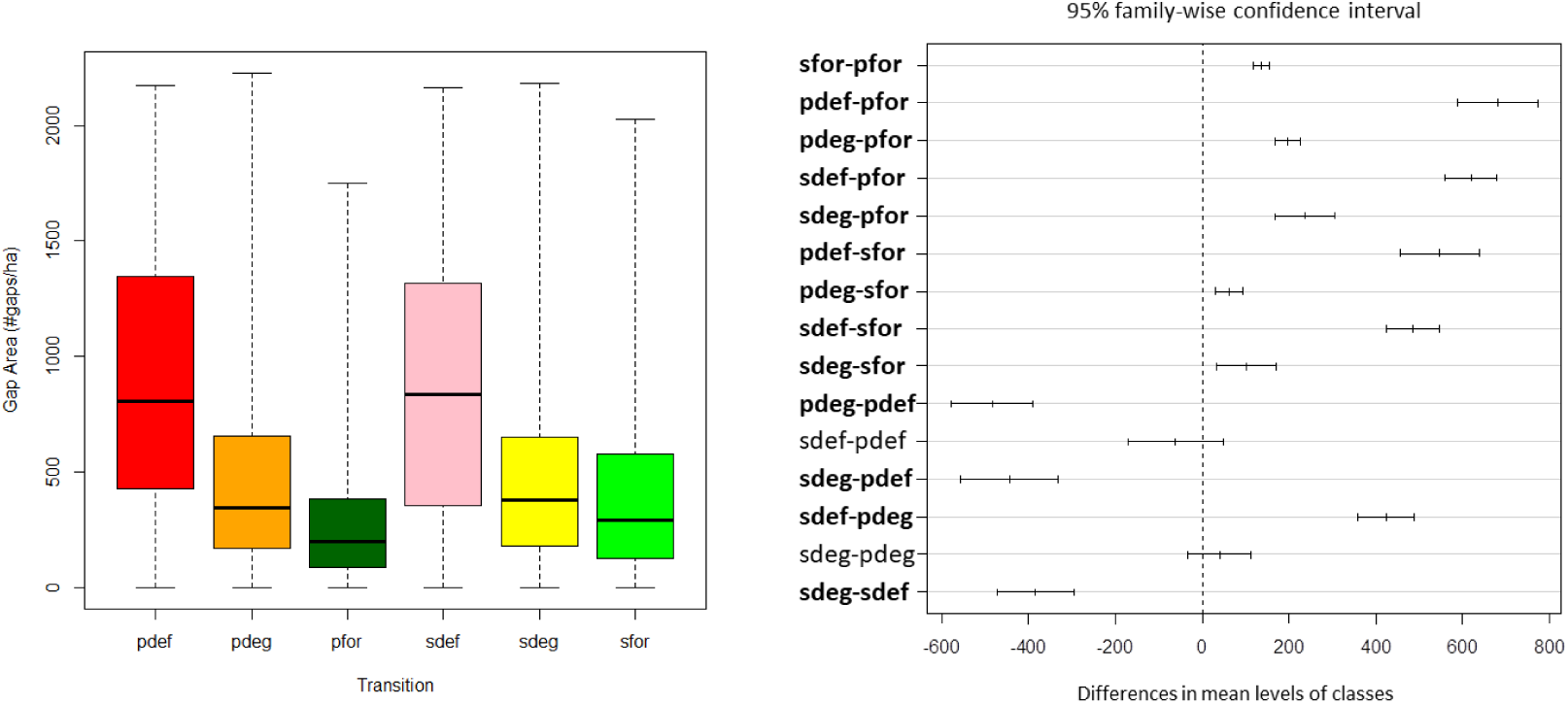
The relationship between gap area per hectare and transition type (left), and Tukey’s HSD comparison of means, (right). Bold indicates significant difference between pairs. The color scheme matches the transitions in figure 2, and from Shapiro et al., 2016. (pdef = primary deforestation; pdeg = primary degradation; pfor = primary forest; sdef = secondary deforestation; sdeg = secondary degradation; sfor = secondary forest)

Mean cumulative negative anomalies were observed to be lowest overall in areas defined as primary or secondary deforestation, and less in degraded areas, and closest to zero in stable forest types with no change. All paired combinations were significantly different, with the exception of primary and secondary deforestation.

### 3.3 Red List of Ecosystems Assessment

Our assessment of the Red List of Ecosystem criteria indicates that 4 ecosystems met criteria to be assessed as critically endangered, 15 as endangered, and 14 as vulnerable (Table 4; Figure 10). The remaining did not meet any of the category thresholds and are therefore listed as least concern. The full table of ecosystems and criteria are presented in the supplementary material, showing that criterion D, which included the forest condition triggered the same red list categories as criterions A and B.

**Table 4.**
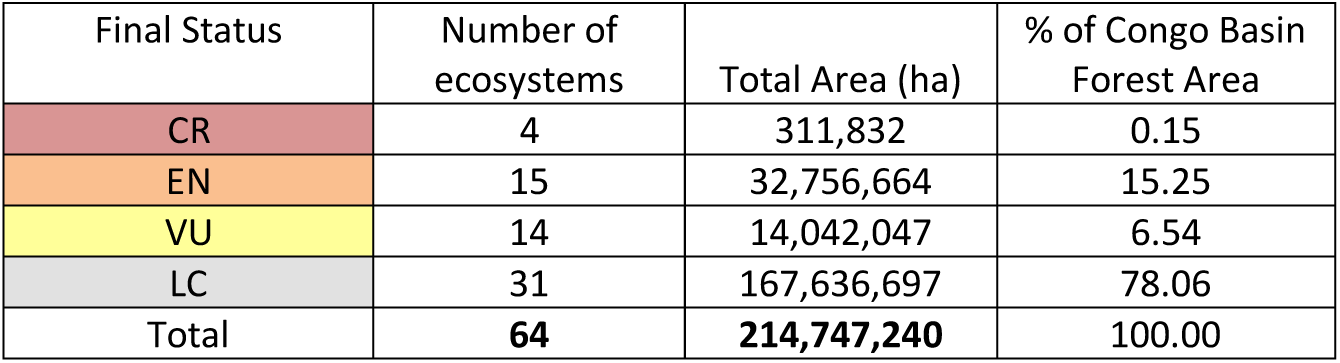
Redlist of Ecosystem summary for 64 Congo Basin Forest Ecosystems

**Figure 9.**
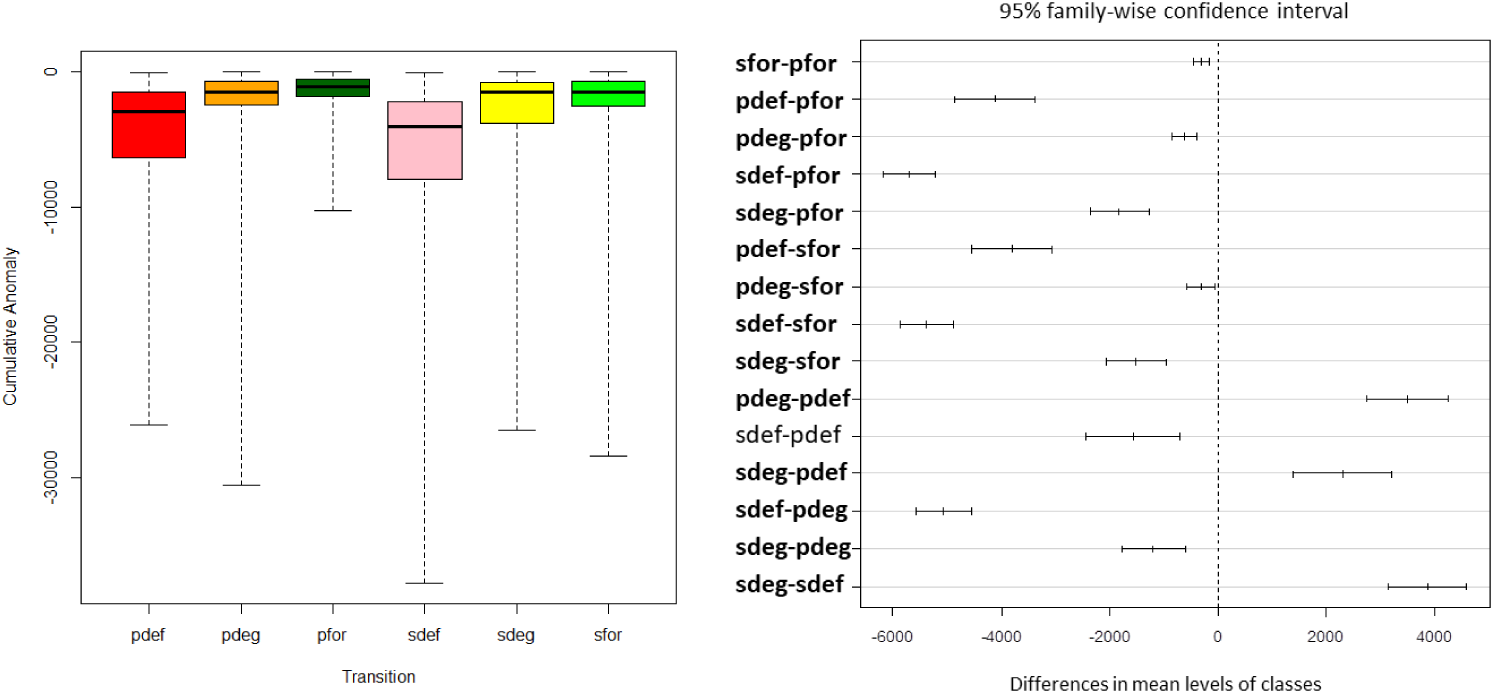
Magnitude of cumulative anomaly by transition type (left), and Tukey’s HSD (right). Bold indicates significant difference between pairs. The color scheme matches the transitions in figure 2, and from Shapiro et al., 2016.

**Figure 10.**
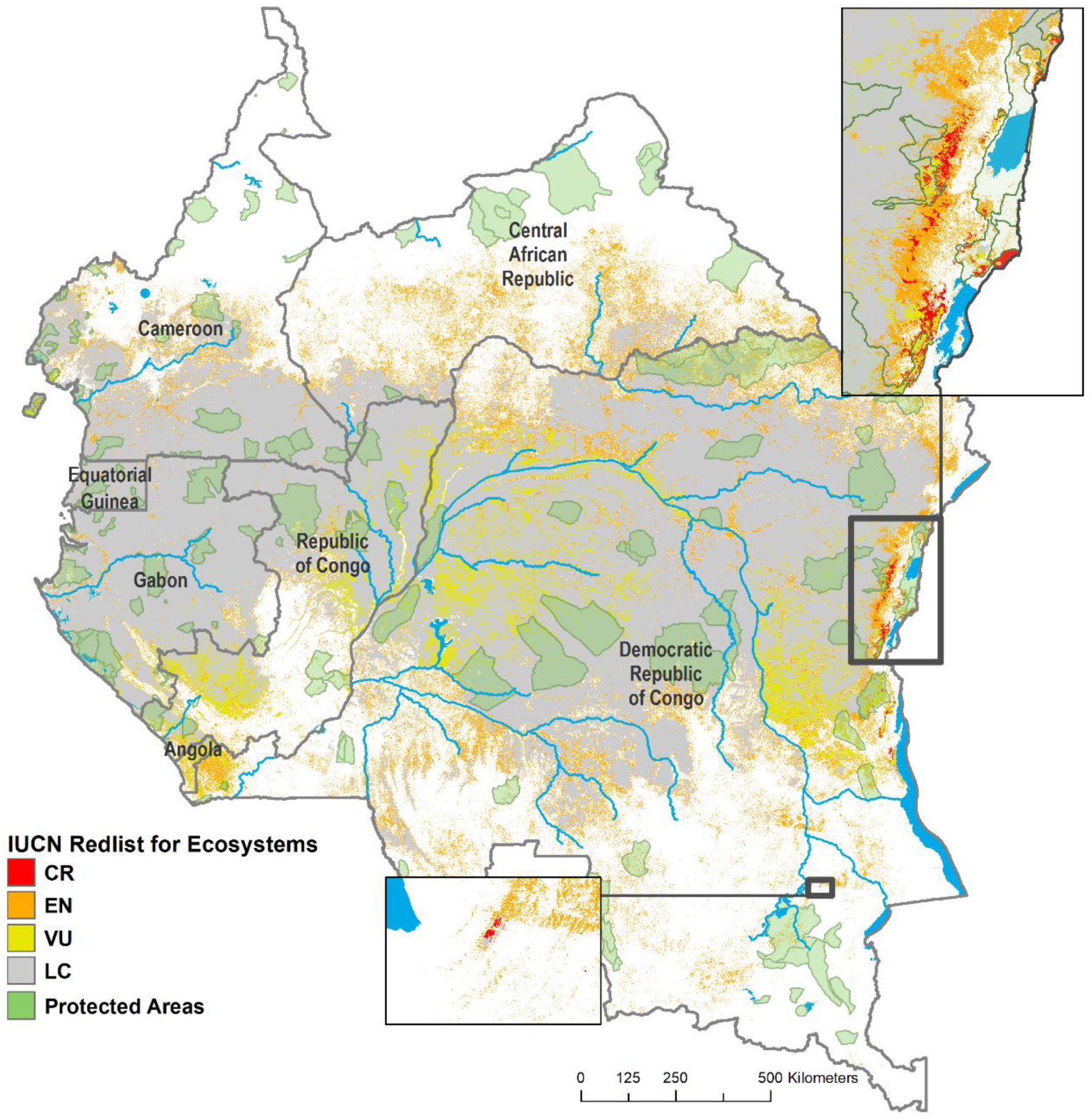
IUCN Redlist assessment for Congo Basin forest ecosystems.

**Figure 11.**
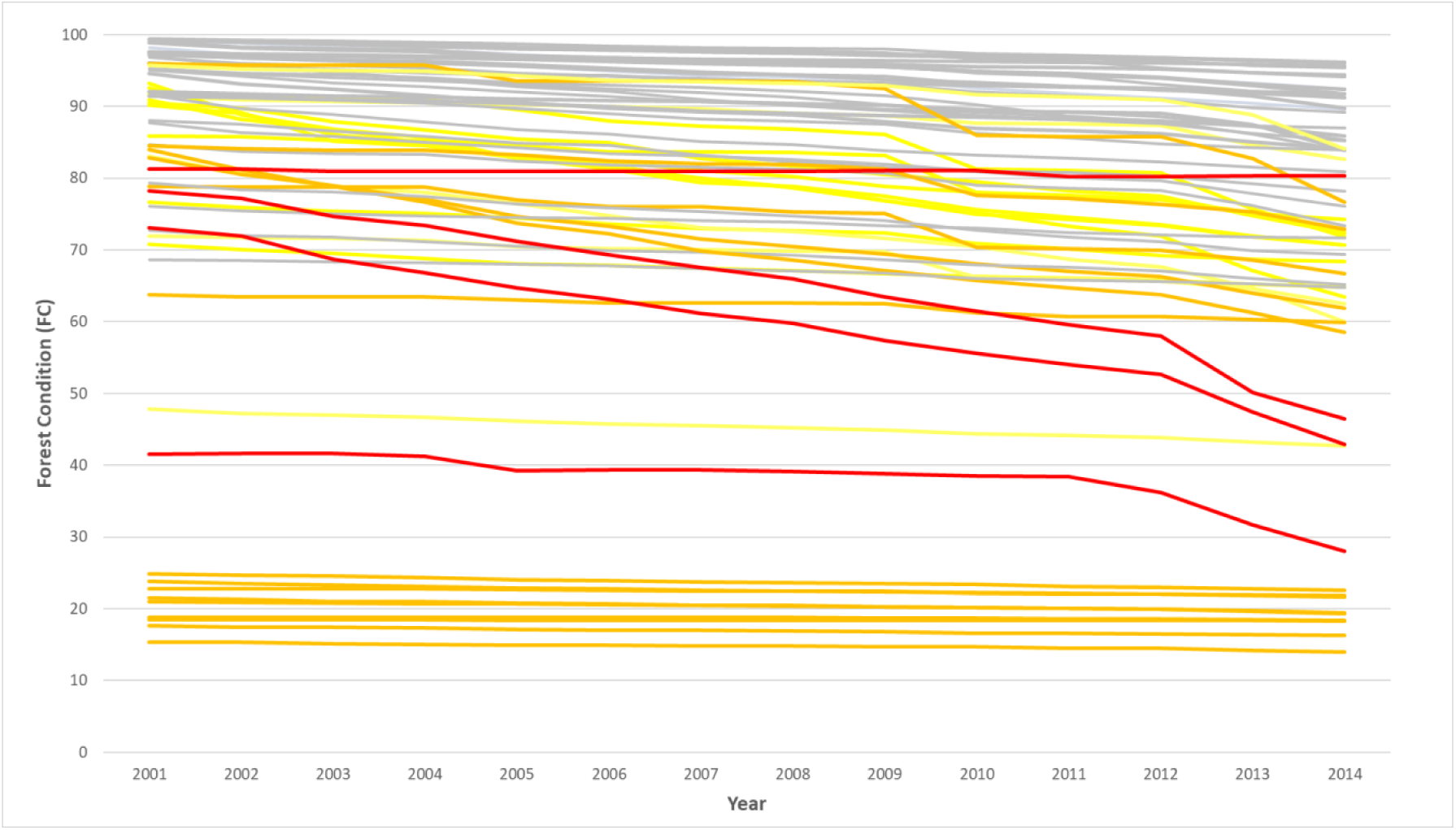
Annual FC of each forest type; grouped into colors according to red list classification (Figure 10).

The four critically endangered ecosystems are located in DRC, notably in and around the Virunga and Kahuzi-Biega National Parks (Figure 10) are shown to have low condition, and experienced significant biomass loss and forest cover loss. DRC also hosts the majority of the endangered ecosystems, along the Congo River and in the west near Angola, along with the northern open forests. Central African Republic is dominated by fragmented, endangered open forests, and the Republic of Congo has large areas of vulnerable ecosystems. In the central cuvette, swamp forests are vulnerable in DRC and Republic of Congo. Several dense, evergreen and semi-deciduous forests in the northeast and northwest regions fall in the endangered categories, while three types of swamp forest ecosystems fall in the vulnerable category.

Of the 33 ecosystems qualified as above least concern, 21 qualified for ranking in a category above Least Concern for criterion A or B as well as D, indicating general agreement between the criteria (Table 5). An additional 12 ecosystems were assigned a threat ranking according to criterion D alone, meaning they did not undergo a significant change in extent, but rather extensive and significantly decreasing condition. These ecosystems included several categories of open forests, which were assigned the higher threat class of endangered due to the extent and severity thresholds for criterion D, while all four critically endangered ecosystems were assigned a higher risk class due to criterion D than A or D. In contrast, no ecosystems were assigned a threat status according to A or B alone, which is expected as reduced area is associated with a reduced core area.

**Table 5.** Ecosystem Redlist assessment for 64 Congo Basin Ecosystems based on criterion A2b, B1 and B2 and D

| Code | General Forest Type | Forest Type<br>Extent in 2000<br>(ha) | Forest Type<br>Extent 2016<br>(ha) | A2b<br>(adjusted %<br>PRD) | B1 Extent of<br>Occurrence<br>(km2) | B2 Area of<br>Occurrence<br>(1% rule) | D2a affected<br>extent<br>(estimated %<br>PRD core forest<br>in 2050) | D2a severity<br>(2050<br>estimated %<br>PRD forest<br>condition) | D3 extent<br>(% core forest<br>loss since<br>1850) | D3 Severity<br>(condition<br>lost since<br>1850) | Final<br>Threat<br>Status |
| --- | --- | --- | --- | --- | --- | --- | --- | --- | --- | --- | --- |
| 11320 | Dense Evergreen | 1,414 | 1,332 | 28.27 | 117,779 | 6 | 87.37 | 90.44 | 84.79 | 71.94 | CR |
| 11340 | Dense Evergreen | 64,235 | 63,176 | 9.08 | 31,391 | 44 | 61.73 | 51.95 | 40.31 | 27.63 | VU |
| 12120 | Dense Evergreen | 127,724 | 115,908 | 39.05 | 1,252,571 | 401 | 48.35 | 48.72 | 64.66 | 35.16 | VU |
| 12130 | Dense Evergreen | 293,166 | 271,207 | 31.97 | 1,226,900 | 939 | 55.08 | 68.77 | 79.15 | 57.34 | VU |
| 12140 | Dense Evergreen | 3,683,258 | 3,652,941 | 3.84 | 1,805,672 | 2523 | 25.36 | 25.99 | 19.49 | 14.09 | LC |
| 12220 | Dense Evergreen | 114,911 | 113,699 | 4.25 | 453,124 | 135 | 20.03 | 18.00 | 11.15 | 8.24 | LC |
| 12240 | Dense Evergreen | 26,543 | 26,321 | 4.70 | 107,017 | 46 | 39.95 | 38.58 | 19.36 | 15.90 | VU |
| 12320 | Dense Evergreen | 185,738 | 180,869 | 11.85 | 167,546 | 121 | 45.80 | 46.44 | 29.46 | 23.93 | LC |
| 12340 | Dense Evergreen | 3,012 | 2,980 | 6.01 | 22,156 | 9 | 48.99 | 54.95 | 41.70 | 33.33 | EN |
| 22120 | Evergreen and Semi-Deciduous | 1,662,006 | 1,650,651 | 3.21 | 1,219,257 | 1318 | 9.13 | 8.85 | 7.31 | 4.33 | LC |
| 22130 | Evergreen and Semi-Deciduous | 385,836 | 369,374 | 19.48 | 1,200,163 | 1059 | 42.08 | 51.35 | 48.55 | 34.78 | LC |
| 22140 | Evergreen and Semi-Deciduous | 14,915,844 | 14,765,946 | 5.04 | 1,886,996 | 5450 | 31.47 | 26.16 | 14.40 | 10.20 | LC |
| 22220 | Evergreen and Semi-Deciduous | 1,029,960 | 1,023,036 | 2.77 | 287,906 | 467 | 15.59 | 11.97 | 5.92 | 3.99 | LC |
| 22240 | Evergreen and Semi-Deciduous | 33,597 | 33,319 | 4.44 | 235,436 | 69 | 36.85 | 33.29 | 23.31 | 16.02 | LC |
| 22320 | Evergreen and Semi-Deciduous | 258,645 | 247,788 | 19.49 | 111,203 | 197 | 60.09 | 58.63 | 33.33 | 27.64 | VU |
| 22340 | Evergreen and Semi-Deciduous | 5,138 | 5,109 | 2.52 | 35,930 | 14 | 31.30 | 48.28 | 50.04 | 40.08 | EN |
| 31110 | Semi-Deciduous | 1,577,743 | 1,549,357 | 7.84 | 193,021 | 701 | 27.20 | 38.61 | 41.60 | 28.35 | LC |
| 31120 | Semi-Deciduous | 29,266,061 | 27,585,817 | 24.46 | 1,279,272 | 5896 | 44.44 | 42.17 | 26.97 | 19.10 | LC |
| 31130 | Semi-Deciduous | 13,376,916 | 12,696,521 | 23.18 | 961,081 | 3664 | 44.45 | 38.95 | 22.86 | 15.88 | LC |
| 31140 | Semi-Deciduous | 37,851,190 | 37,051,712 | 10.24 | 1,924,336 | 7737 | 41.80 | 35.57 | 21.42 | 14.70 | LC |
| 31220 | Semi-Deciduous | 498,705 | 454,093 | 34.18 | 369,112 | 392 | 80.45 | 72.88 | 53.28 | 38.12 | EN |
| 31240 | Semi-Deciduous | 75,450 | 74,388 | 7.09 | 252,923 | 89 | 55.07 | 47.77 | 41.55 | 26.64 | LC |
| 31320 | Semi-Deciduous | 129,090 | 116,523 | 41.88 | 111,092 | 142 | 88.86 | 87.53 | 61.91 | 53.53 | CR |
| 31340 | Semi-Deciduous | 14,949 | 14,806 | 4.96 | 39,374 | 27 | 39.95 | 54.97 | 45.78 | 37.49 | VU |
| 32120 | Semi-Deciduous | 8,740,924 | 8,627,203 | 6.48 | 911,114 | 2573 | 19.21 | 14.07 | 7.64 | 4.69 | LC |
| 32130 | Semi-Deciduous | 10,705,279 | 10,592,702 | 5.23 | 1,166,921 | 4396 | 14.98 | 14.22 | 8.27 | 5.88 | LC |
| 32140 | Semi-Deciduous | 3,190,203 | 3,136,105 | 7.68 | 1,019,159 | 3006 | 44.24 | 38.61 | 22.30 | 16.17 | LC |
| 32220 | Semi-Deciduous | 810,449 | 783,165 | 14.73 | 428,420 | 560 | 48.36 | 39.39 | 21.77 | 14.81 | LC |
| 32240 | Semi-Deciduous | 10,654 | 10,557 | 5.10 | 200,429 | 35 | 47.46 | 36.06 | 29.31 | 17.40 | VU |
| 32320 | Semi-Deciduous | 208,448 | 192,664 | 33.72 | 110,475 | 197 | 76.22 | 73.76 | 43.57 | 36.58 | VU |
| 32340 | Semi-Deciduous | 2,116 | 2,103 | 3.45 | 18,753 | 5 | 49.69 | 48.41 | 36.81 | 27.12 | EN |
| 42120 | Semi-Deciduous with Pioneer | 3,213,222 | 2,949,922 | 34.01 | 489,205 | 1677 | 60.99 | 57.02 | 34.68 | 25.80 | VU |
| 42130 | Semi-Deciduous with Pioneer | 17,198,872 | 16,027,875 | 28.97 | 1,194,308 | 5693 | 43.40 | 43.08 | 30.27 | 21.82 | LC |
| 42140 | Semi-Deciduous with Pioneer | 2,467,033 | 2,342,183 | 22.49 | 869,131 | 1485 | 73.78 | 72.83 | 47.95 | 40.04 | VU |
| 42220 | Semi-Deciduous with Pioneer | 349,164 | 316,814 | 35.93 | 300,419 | 410 | 83.70 | 77.44 | 55.43 | 41.53 | EN |
| 42230 | Semi-Deciduous with Pioneer | 922 | 901 | 12.73 | 10,118 | 1 | 39.35 | 21.75 | 81.24 | 19.68 | CR |
| 42240 | Semi-Deciduous with Pioneer | 6,468 | 6,351 | 9.67 | 41,384 | 20 | 62.28 | 59.80 | 34.52 | 28.19 | VU |
| 42320 | Semi-Deciduous with Pioneer | 180,406 | 160,827 | 45.23 | 167,140 | 208 | 89.20 | 88.59 | 66.03 | 57.11 | CR |
| 42340 | Semi-Deciduous with Pioneer | 2,334 | 2,291 | 9.58 | 6,542 | 3 | 67.44 | 54.46 | 33.85 | 23.30 | EN |
| 51140 | Maranthaceae | 270,382 | 265,991 | 8.00 | 33,354 | 188 | 33.88 | 22.58 | 15.40 | 8.29 | LC |
| 63151 | Swamp Forest | 4,537,018 | 4,068,227 | 42.14 | 376,553 | 2433 | 74.67 | 67.72 | 38.91 | 29.36 | VU |
| 63152 | Swamp Forest | 6,036,564 | 5,945,728 | 7.41 | 369,866 | 2447 | 25.52 | 22.60 | 9.63 | 7.58 | LC |
| 63153 | Swamp Forest | 1,783,496 | 1,753,679 | 8.00 | 380,728 | 1571 | 32.24 | 28.34 | 14.52 | 10.84 | LC |
| 63154 | Swamp Forest | 817,340 | 801,088 | 9.29 | 368,573 | 909 | 49.86 | 50.93 | 39.79 | 30.68 | LC |
| 63155 | Swamp Forest | 1,033,979 | 1,023,902 | 5.11 | 266,923 | 896 | 29.38 | 25.94 | 10.55 | 8.68 | LC |
| 63156 | Swamp Forest | 1,548,171 | 1,540,305 | 2.72 | 266,485 | 824 | 19.38 | 16.73 | 6.97 | 5.55 | LC |
| 63157 | Swamp Forest | 311,196 | 307,660 | 5.92 | 251,245 | 386 | 33.06 | 30.52 | 18.05 | 13.03 | LC |
| 63161 | Swamp Forest | 1,671,255 | 1,498,735 | 41.92 | 480,285 | 1190 | 72.39 | 65.72 | 39.41 | 29.27 | VU |
| 63162 | Swamp Forest | 4,642,848 | 4,541,822 | 10.07 | 490,315 | 1842 | 32.90 | 29.59 | 13.29 | 10.43 | LC |
| 63163 | Swamp Forest | 3,205,778 | 3,164,469 | 5.84 | 490,983 | 1652 | 25.38 | 22.87 | 11.41 | 8.79 | LC |
| 63164 | Swamp Forest | 1,142,687 | 1,124,208 | 7.22 | 457,531 | 959 | 51.62 | 49.48 | 40.53 | 31.67 | VU |
| 63165 | Swamp Forest | 554,484 | 546,680 | 6.44 | 288,024 | 691 | 25.78 | 22.84 | 9.53 | 7.59 | LC |
| 63166 | Swamp Forest | 2,014,147 | 2,003,064 | 2.72 | 292,569 | 870 | 13.08 | 11.65 | 4.73 | 3.81 | LC |
| 63167 | Swamp Forest | 209,992 | 206,923 | 6.59 | 292,902 | 396 | 35.04 | 33.51 | 21.01 | 16.04 | LC |
| 71140 | Mangrove | 404,083 | 401,897 | 2.54 | 224,232 | 174 | 29.77 | 43.69 | 45.02 | 35.29 | LC |
| 81110 | Open Forest | 9,077,498 | 8,912,775 | 8.12 | 1,623,726 | 6243 | 55.48 | 88.32 | 96.92 | 80.60 | EN |
| 81120 | Open Forest | 7,915,813 | 7,287,374 | 31.61 | 1,051,266 | 4280 | 66.17 | 76.39 | 97.38 | 78.21 | EN |
| 81140 | Open Forest | 2,752,083 | 2,585,455 | 26.52 | 2,773,294 | 2570 | 56.22 | 85.19 | 95.00 | 80.74 | EN |
| 81220 | Open Forest | 577,782 | 544,560 | 23.91 | 205,313 | 431 | 64.98 | 82.46 | 97.06 | 77.48 | EN |
| 81240 | Open Forest | 85,797 | 85,004 | 4.39 | 261,394 | 136 | 51.30 | 81.04 | 96.18 | 78.42 | EN |
| 81320 | Open Forest | 282,233 | 263,227 | 28.64 | 80,439 | 229 | 70.74 | 88.77 | 98.32 | 86.00 | EN |
| 81340 | Open Forest | 38,659 | 38,139 | 5.67 | 90,736 | 62 | 30.56 | 82.49 | 95.91 | 81.59 | EN |
| 82130 | Open Forest | 10,614,508 | 9,686,379 | 34.14 | 1,967,561 | 8913 | 70.42 | 86.99 | 98.92 | 83.75 | EN |
| 82230 | Open Forest | 551,822 | 540,422 | 9.61 | 908,869 | 846 | 35.67 | 82.29 | 99.56 | 81.74 | EN |

The trajectory of FC over time assessed differs for each ecosystem and redlist category. The critically endangered ecosystems are shown to decrease more rapidly after 2012, with the exception of the evergreen/semi-deciduous ecosystem (upper most line) which has a slow decline in FC, but it is particularly affected by its limited extent. The lowest lines represent the open forests which overall lower condition compared to other ecosystem types, as they are greatly fragmented and as a result have a much lower than the maximum potential AGB.

## 4. Discussion

Our research identified 64 unique forest ecosystems which provides a basis for representative and comprehensive conservation planning across international boundaries, climate zones and biogeographical regions. Although forest cover is still extensive, the impacts of fragmentation and biomass loss occur throughout the region (33% of overall forest area), reducing the ecological condition of forests, notably in the eastern mountains of DRC and southern, northern peripheries of semi-deciduous forests stands; the open forests of Central African Republic and southern Cameroon. Through our analysis we have developed functional tools to support the RLE by defining ecosystems with reduced extent and significantly reduced condition. The application of FC to evaluate potential ecosystem collapse has provided additional information than extent or size alone (criterions A and B).

We characterize FC as a combination of biomass lost and spatial pattern in time to produce a metric on a continuous scale from 0-100%. In contrast with indicators that provide a single snapshot in time, or binary assessments of intact vs. not (Potapov et al., 2008; Tyukavina et al., 2016), FC has the unique element of incorporating a temporal dimension of biomass loss to produce a relative index of degradation. This is an important requirement for accurately describing forest degradation, and differentiating a regenerating secondary forest, from one which is stable, from one which may have previously been intact (Thompson et al., 2013). The overall approach to developing this metric lies in a specific definition of forest degradation, based AGB which is a function of climate regulating services of forests, an essential ecosystem service for preventing climate change provided by intact, functional forests. Therefore, the assessment of FC over time provides an important metric for monitoring forests capacity to either sequester or emit forest carbon over time, but is not limited to such use as it can be used to prioritize restoration efforts.

FC was positively correlated with forest canopy fractional cover, biomass lost over time, and negatively associated gap area at 1 hectare scale, validating the theoretical model of subsequent states of degradation presented in Figure 2. The assessment of forest transitions (primary and secondary deforestation and degradation) gap area and cumulative anomalies of direct assessments of long-term changes in NBR provide more context in describing the successive forest states which lead to deforestation. The incremental significant differences point to an indicator which can accurately discern deforestation from degradation, and the combination of temporal data with biomass allows for more information than any of these variables alone. FC and transitions together provide an informative stratification for cost-effective conservation planning, monitoring and climate change interventions, as direct measures of forest gaps, fractional cover or direct remote sensing metrics alone do not inform the prior status of a forest ecosystem. High resolution forest structure and gap area require significant investments into very high resolution airborne or drone data which are not always feasible. While fractional cover remains highly correlated with the other validation variables, fractional tree cover from satellite cannot adequately discern different forest heights or high or low biomass ecosystems. Additionally, a forest with a continuous canopy will have the same fractional cover regardless of its biomass, structure, making it inadequate to independently assess relative degradation state.

The Landsat observation frequency is not always ideal for wall-to-wall degradation detection, particularly before Sentinel-2, and higher resolution sensors such as Planet data have cost barriers and are less spectrally consistent than lower resolution sensors. For these reasons, an indirect method, although it might incur greater assumptions and over-simplification of processes, can be more adapted and flexible to quick monitoring assessments. Although indirect methods are generally simpler, they can provide the necessary information for conservation planning, stratification for emissions reduction project reporting (Pelletier et al., 2013). In fact, we showed that the integration of temporal information can differentiate primary from secondary deforestation where a direct spectral measure or estimation of fractional cover cannot.

Long term cumulative NBR anomalies are feasible to estimate with satellite imagery and powerful cloud computing, but alone cannot discern deforestation events which may be followed by quick regeneration, nor does it differentiate between different types of forest dynamics. An assessment of trends, for example using LandTrendr (Kennedy et al., 2010), might be required to investigate various transitions, but still aren’t designed to assess the relative changes occurring in primary or secondary forest types, and these require consistent long term cloud-free time series data and calibration information that remain sensitive to short term dynamics.

Tukey HSD pairings of differences in mean canopy gap, and anomalies are significantly different, with the exception of primary and secondary deforestation, which were not significant in the paired variable tests. This is not entirely unexpected, as a deforested ecosystems are similar whether it was previously intact or already degraded. For this reason, FC provides important contextual information, to differentiate the differences in subsequent degradation transitions from stable secondary or degraded forests and provides a suitable indirect method to meet most monitoring needs.

Our approach of monitoring FC over time is valuable for conservation prioritization and planning or stratification of areas of intervention, notably via the ecosystem redlist. We observed varying FC for ecosystems, where areas with lower condition may be prioritized for restoration activities, while those with high overall condition could be managed for conservation and carbon stock maintenance. In comparison with binary indices such as Intact Forest Landscapes (Potapov et al., 2008), hinterland forests (Tyukavina et al., 2016) or identifying core and edge (N. M. Haddad et al., 2015; Riitters et al., 2015), FC provides a continuous index which can be applied in a more flexible manner for conservation planning and assessment of ecosystem intactness. In particular in the Congo basin, FC quantifies many forests which happen to fall outside the IFL definition yet are the locations of essential corridors, valuable great ape habitat, or are identified as Key Biodiversity Areas (KBAs) (IUCN, 2016b; The KBA Partnership, 2018). In addition, a continuous metric integrating the temporal history supports conservation prioritization and methods to rank areas by FC for different interventions – such as active restoration or mitigation activities to promote regeneration.

### 4.1 Application to IUCN Red List of Ecosystems

The application of FC into Criterion D of the IUCN Red List enables us to assess the disruption of biotic processes over a large region, assessing both spatial extent of impact and the severity, which could be otherwise difficult to measure or estimate, for example biotic processes related to species richness, trophic diversity (see supplementary material of Bland et al., 2015). We found that we would have under-estimated ecosystem risk by assigning the Red List category based only on the geographic extent and area of occupancy by applying criterion A and B only. The additional element of FC is necessary to assess ecosystem status independent of spatial extent. As our analysis has shown, several ecosystems did not meet the extent criteria for A or B and might not be geographically “rare”, but were triggered by criterion D, while no ecosystems were classified at risk with only A or B. This was observed in all open forest categories in dryland ecosystems which despite their very fragmented state can still potentially harbour high AGB (Bastin et al., 2017). This high potential AGB results in low relative FC which triggers criterion D, while their large geographic distribution do not trigger criterion A or B. This shows that A and B do not adequately integrate fragmentation and pattern to assess ecosystems. This also indicates that criterion D captures the ranking of several criteria and is an effective indicator for the ecosystem risk assessment, as it measures both extent and condition, as opposed to A and B which are focused primarily on extent. As ecosystem functioning, notably biodiversity greatly affected by fragmentation (Haddad et al., 2015), it is logical and necessary to include spatial pattern metrics in an ecosystem risk assessment designed for conservation. The FC estimate directly addresses the concept of the endpoint (FC=0) of ecosystem decline, supporting the scientific underpinning of the ecosystem red list process (Keith et al., 2013) and can also be applied to other ecosystem prioritization efforts for conservation. Finally, this assessment has shown that that the principal driver of ecosystem collapse in Congolese forest systems are related to fragmentation and degradation, and while deforestation overall may remain low, there are significant pressures that can affect forest health and associated biodiversity (Grantham et al., 2020).

The availability of temporal data and trends over annual time steps enables a forward and backward modelling to fit the criteria requirements of estimating changes in FC over 50 years past or future predictions. Most importantly, the method has enabled the identification of critically endangered ecosystems among the large extensive forests in the Congo Basin. In particular, the montane and sub-montane forests identified as critically endangered are already limited in extent and have suffered deforestation and degradation, and are home to the Eastern Chimpanzee and Eastern Gorilla habitat, which are endangered and critically endangered species on the IUCN Red List respectively (IUCN, 2019). These habitats are presently within iconic protected areas such as Virungas National Park, which have undergone recent forest loss and threats from oil development, demonstrating the limits of formal protection and World Heritage status in a situation of political instability, high levels of poverty, and conflict (Hochleithner, 2017; United Nations Economic Commission for Africa, 2015). The other critically endangered habitat identified is currently unprotected and lies between several mining concessions which might present acute threats in the future (Pélissier et al., 2019). Additionally, particular consideration should be given to ecosystems in endangered and vulnerable categories, which lie along southern edge of the dense forest ranges. These are more accessible by humans and are present among mixed agricultural landscapes and could be sites to focus restoration activities.

### 4.2 Limitations

All metrics or approximations such as indirect methods or proxies come with the risk of oversimplifying or missing crucial detail that one might observe with direct methods, or for example observing forest degradation events with very high-resolution imagery. FC relies on accurate forest cover maps, which are not always possible with limited validation or available quality data, or at regional scales can be affected by varying forest definitions. For example, the global forest cover maps from Hansen et al., (2013), which are most often used due to access, consistency, resolution, can be difficult to harmonize at the regional global scale because the forest cover threshold varies by latitude, along with different country definitions of forest (Romijn et al., 2013). For this reason, we developed forest ecosystem maps integrating data from various sources and validated with expert opinion to limit bias from one dataset.

FC is a relative index based on biomass estimations, which will always include an element of uncertainty. We overcome this by not using AGB data directly, but rather averaged over forest strata, which should minimize any large errors or inconsistencies, unless most of the ecosystem is already degraded. Additionally, as the changes in biomass are relative, the actual biomass estimates do not necessarily bias the final condition estimate – if biomass is generally over or under-estimated the condition value is not affected. Next, the estimate of maximum potential FC depends on the biomass of forest types at an initial, presumably intact state. For forest types which are already degraded or have low biomass initially based, subsequent condition estimates will be related. For this reason, we recommend that the forest condition index in tandem with the transition classes to adequately identify the current state in the potential degradation time series.

### 4.3 Future Work/implications

Detecting changes in forest cover condition and degradation alone does not meet all the needs for management in the face of increasing population and threats, and new drivers of changing climate. A further step in the analysis is to undertake a geo-spatial assessment of drivers of deforestation and degradation, or their resulting land uses. An assessment of shifting cultivation drivers and change is provided by Molinario *et al*. (2015) which adds a further relevant level of refinement to assign types of transitions to drivers or assess post deforestation land covers. A more in-depth analysis of the complex interactions and changes in drivers over time could provide a finer assessment to manage the causes of deforestation in DRC and define and project future risks and scenario assessment.

## 5 Conclusions

The outlined approach to assessing FC provides a consistent and repeatable tool for evaluating forest over time allowing to distinguish stable, degenerating and regenerating forest. Such distinction is important for the effective orientation of conservation and restoration efforts.. The metric is simple to implement and understand and can provide more information than binary or categorical assessments. We showed that for understanding the threatened status of ecosystems, that understanding their condition can be just as important as its change in extent or rarity.

## Acknowledgements

This work was initiated and coordinated by the Congo basin office of the Forest Stewardship Council (FSC) in the context of their High Conservation Value Roadmap project funded by the Central African Forests Commission (COMIFAC) through its Programme for the Promotion of Certified Logging (PPECF). The authors wish to thank the Forest Stewardship Council for having created the arena for the collaboration between CIRAD, WCS, WWF Germany and WRI. We would also like to express our thanks to WWF-US who supported the organization of the key workshop that brought the main authors of this work together.

## Author contributions

A.S., H.G., V.G., D.B., O.R. developed the forest ecosystem map. A.S. and H.G. developed the forest condition metric, N.A.A. supported A.S. for the biomass and fragmentation assessments; A.S., H.G., N.M and O.R. evaluated the ecosystem data for the risk assessments; O.R. led the project design, coordination and funding acquisition.

