## Supplementary Material for "Forest condition in the Congo Basin for the assessment of ecosystem conservation status"

Table S1. Forest Ecosystem Types of the Congo Basin are coded according to 5 groupings and data sources.

| **Broad ecosystem type** | **Code** |
|---|---|
| Evergreen Rainforest (from Philippon et al., 2018) | 10000 |
| Evergreen and semi-deciduous Rainforest (from Philippon et al., 2018) | 20000 |
| Semi-deciduous Rainforest (from Philippon et al., 2018) | 30000 |
| Semi-deciduous Rainforest with pioneer (from Philippon et al., 2018) | 40000 |
| Marantaceae (zone defined from expert input) | 50000 |
| Swamp Forest (from Betbeder et al., 2014) | 60000 |
| Mangrove (from Giri et al., 2011) | 70000 |
| Open forest (from Hansen et al., 2013) | 80000 |
| **Climate** | |
| northern equatorial (from Philippon et al., 2018) | 1000 |
| southern equatorial (from Philippon et al., 2018) | 2000 |
| central (for swamp forests, from Betbeder et al., 2014) | 3000 |
| **Elevation** | |
| lowland (0-1100m above sea level) | 100 |
| submontane (1100-1750m) | 200 |
| montane (>1750m) | 300 |
| **Biogeographical** | |
| northern (north of Equator) | 10 |
| northern eastern - North and east of Congo River | 20 |
| southern - south of Congo River | 30 |
| north western - north and west of Ubangi river | 40 |
| eastern - east of Congo River | 50 |
| western - West of Ubangi and Congo rivers | 60 |
| **Flooding and swamp forests** | |
| Irregularly Flooded Swamp Forest (from Betbeder et al., 2014) | 1 |
| Seasonal Short-Lasting Flood Pulse Swamp (from Betbeder et al., 2014) | 2 |
| Stable Water Level Swamp Forest (from Betbeder et al., 2014) | 3 |
| Seasonal Flood Pulse Swamp Forest (from Betbeder et al., 2014) | 4 |
| Palm-Dominated Seasonal Short-Lasting Flood Pulse Swamp Forest (from Betbeder et al., 2014 and Dargie et al., 2017) | 5 |
| Palm-Dominated Stable Water Level Swamp Forest (from Betbeder et al., 2014 and Dargie et al., 2017) | 6 |
| Palm-Dominated Seasonal Flood Pulse Swamp Forest (from Betbeder et al., 2014 and Dargie et al., 2017) | 7 |
